# Planarian stem cells sense the identity of missing tissues to launch targeted regeneration

**DOI:** 10.1101/2020.05.05.077875

**Authors:** Tisha E. Bohr, Divya A. Shiroor, Carolyn E. Adler

## Abstract

In order to regenerate tissues successfully, stem cells must first detect injuries and then produce missing cell types through largely unknown mechanisms. Planarian flatworms have an extensive stem cell population responsible for regenerating any organ after amputation. Here, we compare stem cell responses to different injuries by amputation of a single organ, the pharynx, or removal of tissues from other organs by decapitation. We find that planarian stem cells adopt distinct behaviors depending on what tissue is missing: loss of non-pharyngeal tissues increases numbers of non-pharyngeal progenitors, while removal of the pharynx specifically triggers proliferation and expansion of pharynx progenitors. By pharmacologically inhibiting either proliferation or activation of the MAP kinase ERK, we identify a narrow window of time during which proliferation, followed by ERK signaling, produces pharynx progenitors necessary for regeneration. Further, unlike pharynx regeneration, eye regeneration does not depend on proliferation or ERK activation. These results indicate that stem cells tailor their proliferation and expansion to match the regenerative needs of the animal.

## Introduction

When faced with injury or disease, many animals can repair or even replace damaged tissue. This process of regeneration is observed across animal species, and is often fueled by tissue-resident stem cells (Bely & Nyberg, 2010; Alejandro Sánchez Alvarado & Tsonis, 2006; Tanaka & Reddien, 2011). In response to injury, stem cells accelerate the production of specific types of differentiated cells in order to repair damaged tissues. For example, in adult mammals, injuries to the intestine, skin or lung induce stem cells to increase proliferation rates and alter their differentiation potential (Buczacki et al., 2013; Stabler & Morrisey, 2017; Tetteh et al., 2015; Tumbar et al., 2004). These findings suggest that injury can modify the behavior of stem cells to promote repair, but how these changes contribute to tissue regeneration remains unclear.

The freshwater planarian *Schmidtea mediterranea* is an ideal model organism to study how injury invokes repair due to their virtually endless ability to regenerate (Ivankovic et al., 2019). This ability is driven by an abundant, heterogeneous population of stem cells (Adler & Sánchez Alvarado, 2015; Reddien, 2018; Zhu & Pearson, 2016). Defined by ubiquitous expression of the argonaute transcript *piwi-1* (Reddien et al., 2005), planarian stem cells consist of both pluripotent stem cells (PSCs) capable of reconstituting the entire animal (Wagner et al., 2011) and organ-specific progenitors (Figure 1A) (Scimone, Kravarik, et al., 2014; van Wolfswinkel et al., 2014; Zeng et al., 2018). Organ-specific progenitors contribute to the maintenance and regeneration of planarian organs, including a pharynx, primitive eyes, muscle, intestine, an excretory system and a central nervous system (Figure 1A), all enveloped in epithelium (Roberts-Galbraith & Newmark, 2015). During tissue homeostasis, stem cells replenish these organs by constant turnover (Pellettieri & Sánchez Alvarado, 2007). However, once an injury occurs, stem cells immediately begin to proliferate, and vast transcriptional changes ensue (Sandmann et al., 2011; Wenemoser et al., 2012; Wenemoser & Reddien, 2010). Phosphorylation of the extracellular signal-regulated kinase (ERK) is required for many of these wound-induced transcriptional changes, as well as stem cell differentiation, proliferation and survival (Owlarn et al., 2017; Shiroor et al., 2019; Tasaki et al., 2011). How these injury-induced changes regulate stem cell behavior to facilitate the transition from a homeostatic state and contribute to successful regeneration of particular organs are key issues to resolve.

**Figure 1:**
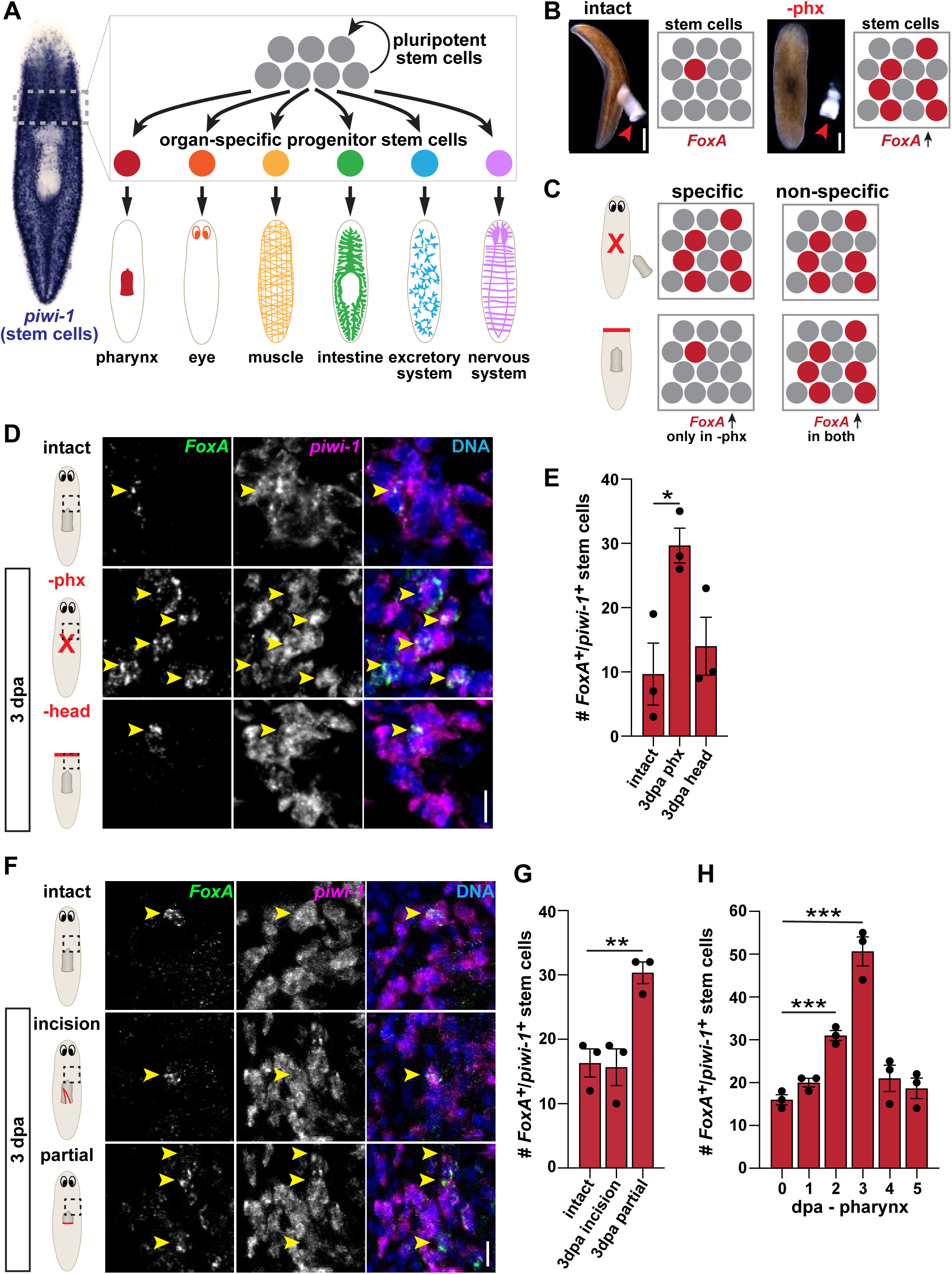
Pharyngeal Progenitors Increase Only after Pharynx Loss. (A) Schematic of planarian stem cell lineage. Left, whole-mount *in situ* hybridization (WISH) for the stem cell marker *piwi-1*. Right, cartoon depiction of the dashed boxed region showing that planarian stem cells consist of both pluripotent stem cells (PSCs) and organ-specific progenitors that produce planarian organs. (B) Live images of planarians before and after pharynx amputation. Boxes represent schematics of *FoxA*^*+*^ progenitor stem cells (red) among other stem cells (grey), before and after pharynx amputation. Red arrows = pharynx; scale bars = 500μm. (C) Models for specific and non-specific stem cell responses after different amputations (indicated by red lines). *FoxA*^*+*^ progenitor stem cells (red), other stem cells (grey). (D) Confocal images of double fluorescent *in situ* hybridization (FISH) for *FoxA* (green) and *piwi-1* (magenta) in intact animals, or 3 days after pharynx or head amputation. DNA = DAPI (blue); dashed boxes = region imaged; arrows = double-positive cells; scale bar = 10μm. (E) Graph of *FoxA*^*+*^ *piwi-1*^*+*^ cells in the area outlined by dashed boxes in D. (F) Confocal images of FISH for *FoxA* (green) and *piwi-1* (magenta) in intact animals, or 3 days after incision or partial amputation of the pharynx. DNA = DAPI (blue); red lines = site of incision or tissue removal; dashed box = region imaged; arrows indicate double-positive cells; scale bar = 10µm. (G) Graph of *FoxA*^*+*^ *piwi-1*^*+*^ cells in the area outlined by dashed boxes in E. (H) Graph of *FoxA*^*+*^ *piwi-1*^*+*^ cells at indicated times after pharynx amputation in the area outlined by dashed boxes in E. Graphs are represented as mean ± SEM. Dots = individual animals; and *, p ≤ 0.05; **, p ≤ 0.01; ***, p ≤ 0.001, unpaired t-test.

Recent transcriptional profiling of planarian stem cells (Fincher et al., 2018; Plass et al., 2018; Scimone, Kravarik, et al., 2014; van Wolfswinkel et al., 2014; Zeng et al., 2018) has revealed markers of organ-specific progenitors, providing an opportunity to track how stem cells respond to injuries and initiate organ regeneration. Because most planarian organs are dispersed throughout the body (Figure 1A), injuries often cause simultaneous damage to multiple organs (Elliott & Sánchez Alvarado, 2013). The resulting complex regenerative response has limited our ability to decipher how stem cells respond to damage of particular organs. Unlike most other planarian organs, the pharynx is anatomically distinct (Adler & Sánchez Alvarado, 2015; Kreshchenko, 2009). Importantly, it can be completely and selectively removed without perturbing other tissues by brief exposure to sodium azide (Adler et al., 2014; Shiroor et al., 2018). Because only a single organ is removed, pharynx amputation vastly simplifies the regeneration challenge posed to the animal. Additionally, a pharynx-specific progenitor, the Forkhead transcription factor *FoxA*, is expressed in stem cells and increases upon pharynx loss (Adler et al., 2014). These properties allow us to dissect how stem cells respond to loss of a specific organ and are regulated to restore it.

By inflicting different types of injuries to both the pharynx and body, and then tracking organ-specific progenitors, we show that planarian stem cells exhibit distinct responses to loss of different organs. We find that amputation of non-pharyngeal tissues only affects the behavior of non-pharyngeal progenitors. Conversely, removal of pharynx tissue specifically triggers an increase in pharynx progenitors, but not other progenitor types. This increase in pharynx progenitors, and subsequent pharynx regeneration, depends on an initial burst of stem cell proliferation, followed by ERK signaling. Unlike the pharynx, eye regeneration is not dependent on proliferation or ERK signaling. Therefore, we propose that loss of different tissues stimulates unique behaviors in stem cells that channel their outputs towards replacement of missing organs.

## Results

### Pharynx Progenitors Increase Only after Pharynx Loss

Previous work identified the forkhead transcription factor *FoxA* as an essential regulator of pharynx regeneration in planarians (Adler et al., 2014; Scimone, Kravarik, et al., 2014). When the pharynx is present during homeostasis, *FoxA* is expressed in a subset of stem cells. However, upon pharynx removal, the number of stem cells expressing *FoxA* increases (Figure 1B) (Adler et al., 2014). These findings support a model in which pharynx loss stimulates stem cells to initiate pharynx regeneration by selectively upregulating expression of *FoxA*. If this is the case, injuries that do not remove pharynx tissue, like head amputations, should not cause an increase in pharynx progenitors (Figure 1C). However, another study has suggested that stem cells may non-specifically modify their progenitor output based on the size and position of a wound, instead of the identity of missing tissues (LoCascio et al., 2017). If this model is true, then head amputation, where pharyngeal tissue is not removed, should also stimulate an increase in pharynx progenitors (Figure 1C).

To test these two possibilities, we labeled pharynx progenitors with double fluorescent *in situ* hybridization (FISH) for *FoxA* and the stem cell marker *piwi-1* 3 days after pharynx or head amputation. We then quantified pharynx progenitors in a region anterior to the pharynx that was adjacent to wounds left by either head and pharynx amputations. As previously reported, we found that pharynx removal caused a significant increase in pharynx progenitors as compared to intact controls (Figure 1D and 1E) (Adler et al., 2014). By contrast, head amputation did not influence the number of pharynx progenitors, which were similar to intact animals (Figure 1D and 1E). These data indicate that pharynx progenitors are produced in response to pharynx loss, but not injury to other tissue types.

To examine if this response is dependent on organ loss, we inflicted injuries to the pharynx that either removed part of it (∼50-80%), or damaged it without removing any tissue. We then labeled and quantified pharynx progenitors 3 days later. If the pharynx was damaged without tissue removal, no increase in pharynx progenitors occurred. However, if part of the pharynx was removed, we observed a significant increase in pharynx progenitors (Figure 1F and 1G). Therefore, stimulation of pharynx progenitor production requires recognition of lost pharynx tissue, but not necessarily loss of the entire organ.

To determine when pharynx progenitors emerge and how long they persist, we quantified the number of pharynx progenitors at various times after pharynx amputation. Pharynx progenitors emerged within 2 days, peaked at 3 days, and returned to homeostatic levels 4 days after amputation (Figure 1H). Together, these data indicate that planarian stem cells sense the loss of missing pharynx tissue to initiate its specific regeneration through the production of pharynx progenitors.

### Proliferation of *FoxA*^*+*^ Stem Cells is Specifically Triggered by Pharynx Loss

Within hours of any injury, planarian stem cells increase proliferation rates. If tissue is removed, a later, “regeneration-specific” wave of proliferation occurs adjacent to the injury, and persists for days (Baguñà, 1976; Wenemoser & Reddien, 2010). The precision of pharynx amputation, paired with stem cell expression of the pharynx-specific progenitor marker *FoxA* allows us to define the potential contributions of these proliferative cells to the regeneration of a specific organ. By visualizing mitotic figures with antibody staining for phosphohistone H3 (H3P), we confirmed previous results that local stem cell proliferation increases near the wound 1 day after either pharynx or head removal (Figure 2A) (Adler et al., 2014; Baguñà, 1976). To determine if these proliferating stem cells express *FoxA*, we combined FISH for *FoxA* with antibody staining for H3P, 1 day after amputation. Following pharynx removal, we observed higher numbers of *FoxA*^*+*^ H3P^+^ stem cells as compared to intact animals (Figure 2B). To determine how soon after amputation these stem cells initiate proliferation, and how long it persists, we monitored the coincidence of *FoxA*^*+*^ H3P^+^ stem cells at various times after pharynx amputation. Proliferation of *FoxA*^*+*^ stem cells increased within 6 hours of amputation, peaked 1-2 days later, and returned to homeostatic levels 4 days after amputation (Figure 2C). Despite an increase in stem cell proliferation 1 day after head amputation (Figure 2A) (Baguñà, 1976), numbers of *FoxA*^*+*^ H3P^+^ stem cells did not correspondingly increase (Figure 2B and 2D). These data show that pharynx loss specifically stimulates *FoxA*^*+*^ stem cells to proliferate prior to the observed increase in pharynx progenitors that occurs 3 days after amputation (Figure 1H) (Adler et al., 2014).

**Figure 2:**
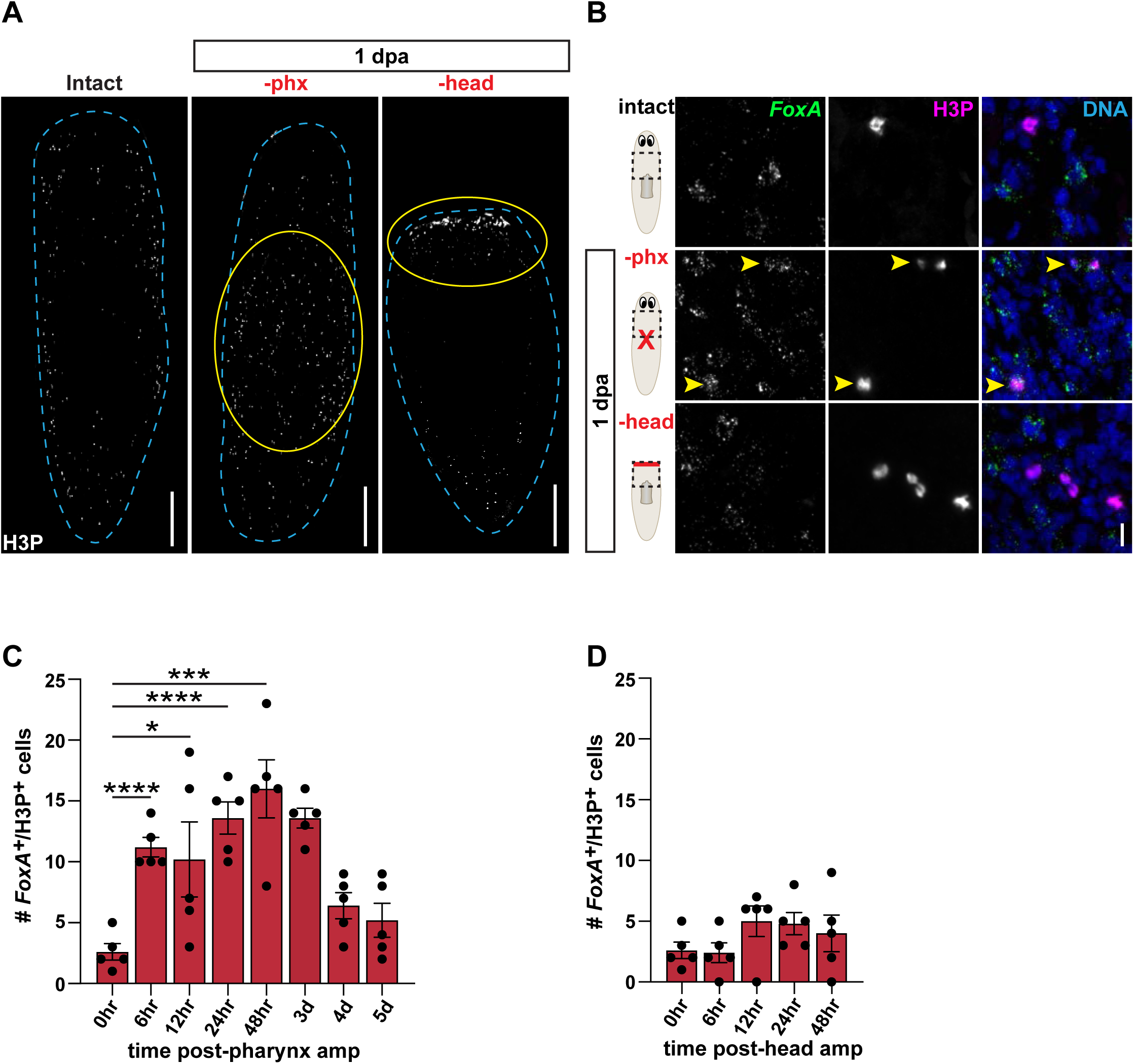
Proliferation of *FoxA*^*+*^ Stem Cells is Specifically Triggered by Pharynx Loss. (A) Whole-mount images of phosphohistone H3 (H3P) antibody in intact worms, or 1 day after pharynx or head amputation. Dashed blue outlines animal; scale bars = 250μm. (B) Confocal images of FISH for *FoxA* (green) and H3P antibody (magenta) in intact animals, or 1 day after pharynx or head amputation. DNA = DAPI (blue); dashed boxes = region imaged; arrows = double-positive cells; scale bar = 10μm. (C) Graph of *FoxA*^*+*^ H3P^+^ cells at indicated times after pharynx amputation in the area outlined by dashed boxes in B. (D) Graph of *FoxA*^*+*^ H3P^+^ cells at indicated times after head amputation in the area outlined by dashed boxes in B. Graphs are represented as mean ± SEM. Dots = individual animals; and *, p ≤ 0.05; ***, p ≤ 0.001, ****, p ≤ 0.0001, unpaired t-test.

Planarian pluripotent stem cells (PSCs) are able to self-renew and generate all the cell types in the body (Figure 1A) (Scimone, Kravarik, et al., 2014; van Wolfswinkel et al., 2014; Wagner et al., 2011; Zeng et al., 2018). The recent identification of molecular markers of PSCs enables tracking of these cells after injuries. One such marker is the tetraspanin group specific gene 1 (*tgs-1*) *(Zeng et al., 2018)*. To identify whether PSCs expressing *FoxA* proliferate after pharynx loss, we labeled mitotic figures with H3P and double FISH for *FoxA* and *tgs-1* 1 day after amputation. Following pharynx amputation, we observed an increase in proliferation of PSCs negative for *FoxA* (*FoxA*^*-*^ *tgs-1*^*+*^), pharynx progenitors (*FoxA*^*+*^ *tgs-1*^*-*^), and PSCs expressing *FoxA* (*FoxA*^*+*^ *tgs-1*^*+*^) (Figure S1A and S1B). Therefore, both *FoxA*^*+*^ PSCs and pharynx progenitors are triggered to divide after pharynx loss.

### Proliferation in a Critical Window of Time is Required for Pharynx Regeneration

In planarians, lineage tracing experiments have shown that proliferation contributes to the production of regenerated tissues (Cowles et al., 2013; Eisenhoffer et al., 2008; Forsthoefel et al., 2011; Newmark & Sánchez Alvarado, 2000; Wagner et al., 2011). We find that proliferation of *FoxA*^*+*^ stem cells (Figure 2C) precedes the increase in pharynx progenitors (Figure 1H) following pharynx amputation, suggesting that stem cell proliferation may generate pharynx progenitors. To test this possibility, we blocked cell division with nocodazole, a microtubule destabilizer that causes a metaphase arrest. In planarians, exposure to nocodazole for 24 hours causes an accumulation of mitotic (H3P^+^) nuclei (Figure 2A) (Grohme et al., 2018; van Wolfswinkel et al., 2014).

We soaked animals in nocodazole for 24 hour increments beginning either immediately after amputation, or 1 or 2 days after pharynx amputation (Figure 3B). Because the pharynx is required to ingest food, amputation causes an inability to eat for about 7 days, until the organ is functionally regenerated (Adler et al., 2014; Ito et al., 2001). Therefore, to determine the outcome of this transient blockade of mitosis on pharynx regeneration, we assayed the recovery of feeding behavior. Animals treated with nocodazole from 0-1 or 2-3 days after pharynx amputation recovered feeding behavior at a similar rate as DMSO-treated control animals (Figure 3C). However, animals treated with nocodazole 1-2 days after pharynx amputation had a drastic delay in recovery of feeding, with only 50% of worms regaining the ability to eat within 20 days of amputation, and 100% within 32 days of amputation (Figure 3C). To verify that nocodazole treatment under these conditions delayed pharynx regeneration, we examined pharynx anatomy 7 days after amputation by whole-mount FISH of the pharynx-specific marker *laminin (Cebrià et al., 2007)*. Animals treated with nocodazole 1-2 days after pharynx amputation had markedly smaller pharynges as compared to DMSO-treated controls (Figure 3D). Because such a transient inhibition of proliferation significantly delayed pharynx regeneration, we conclude that stem cell division 1-2 days after amputation is critical for pharynx regeneration.

**Figure 3:**
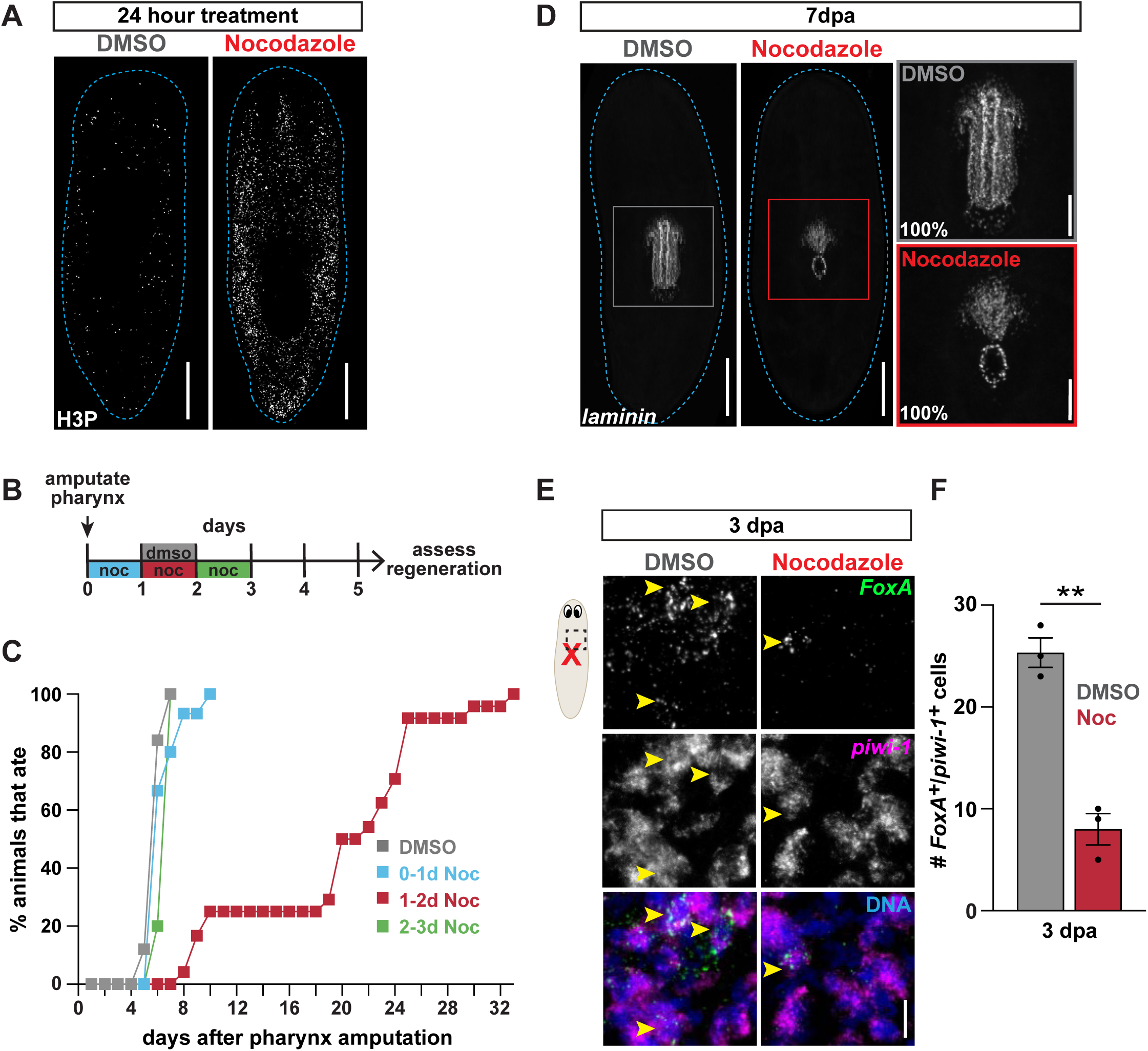
Proliferation in a Critical Window of Time is Required for Pharynx Regeneration. (A) Whole-mount images of H3P antibody in intact animals treated with DMSO (control) or nocodazole for 24 hours. Dashed line outlines animal; scale bars = 250µm. (B) Schematic of nocodazole treatment relative to pharynx amputation for graph in C. (C) Graph of feeding behavior of animals after pharynx amputation, treated as indicated in B and assayed daily, represented as a proportion. n ≥ 20 animals. (D) Whole-mount FISH for the pharynx marker *laminin* 7 days after pharynx amputation in animals treated with DMSO or nocodazole, 1 day after amputation for 24 hours. Dashed line outlines animal; boxes = zoomed area on right; scale bars = 250µm (left), 50µm (right). n ≥ 10 animals. (E) Confocal images of FISH for *FoxA* (green) and *piwi-1* (magenta) 3 days after pharynx amputation in animals treated with DMSO or nocodazole, 1 day after amputation for 24 hours. DNA = DAPI (blue); dashed boxes = region imaged; arrows = double-positive cells; scale bar = 10µm. (F) Graph of *FoxA*^*+*^ *piwi-1*^*+*^ cells in the area outlined by dashed boxes in E, represented as mean ± SEM. Dots = individual animals; and **, p ≤ 0.01, unpaired t-test.

Notably, this time frame coincides with the proliferative peak of *FoxA*^*+*^ stem cells that occurs after pharynx amputation (Figure 2C), suggesting that proliferation directly contributes to the increase in pharynx progenitors observed 3 days after amputation (Figure 1H). To test whether proliferation in this brief window is required for the production of pharynx progenitors, we analyzed the impact of nocodazole exposure on *FoxA*^*+*^ stem cells. First, we examined whether *FoxA*^*+*^ stem cells were mitotically arrested following nocodazole treatment. We observed a significant increase in *FoxA*^*+*^ H3P^+^ stem cells 2 days after amputation (Figure S2A and S2B), illustrating that stem cells poised to become pharyngeal cells were arrested in mitosis. We then performed FISH for *FoxA* and *piwi-1* 3 days after pharynx amputation and found that nocodazole treatment caused a dramatic decrease in pharynx progenitors compared to DMSO-treated controls (Figure 3E and 3F). Importantly, intact animals treated with a similar exposure to nocodazole for 24 hours followed by 1 day of recovery showed no difference in the abundance of pharynx progenitors compared to DMSO-treated controls (Figure S2C and S2D), indicating that proliferation in this short time frame affects pharynx progenitors specifically during regeneration. Together, our data show that proliferation in a critical window of 1 to 2 days after pharynx amputation produces a population of progenitors that are likely essential for pharynx regeneration.

### ERK Phosphorylation is Required for Increased Pharyngeal Progenitors

The mitogen activated protein (MAP) kinase pathway drives proliferation and differentiation during development and regeneration in many organisms (Ghilardi et al., 2020; Patel & Shvartsman, 2018). In planarians, the MAP kinase ERK is the earliest known signal required for regeneration. ERK is phosphorylated within minutes after an injury, which is dispensable for wound healing but required for regeneration and stem cell proliferation (Owlarn et al., 2017; Tasaki et al., 2011). To determine whether ERK is required for pharynx regeneration, we exposed animals to PD0325901 (PD), an inhibitor of the upstream kinase MEK which normally phosphorylates ERK. Following pharynx amputation, we maintained animals in the presence of PD for 5 days (Figure 4A), which was previously shown to inhibit head regeneration (Owlarn et al., 2017), and then assayed feeding behavior daily. While control animals regained the ability to feed within 7 days, animals treated with PD had a substantial delay in feeding, with 50% of worms feeding by day 13 and all worms feeding by day 29 (Figure 4B). We verified that this delay in feeding was due to defects in pharynx regeneration with whole-mount *laminin* FISH, and found that PD-treated animals lacked a pharynx 7 days after amputation (Figure 4C). These data indicate that ERK signaling is required for pharynx regeneration.

**Figure 4:**
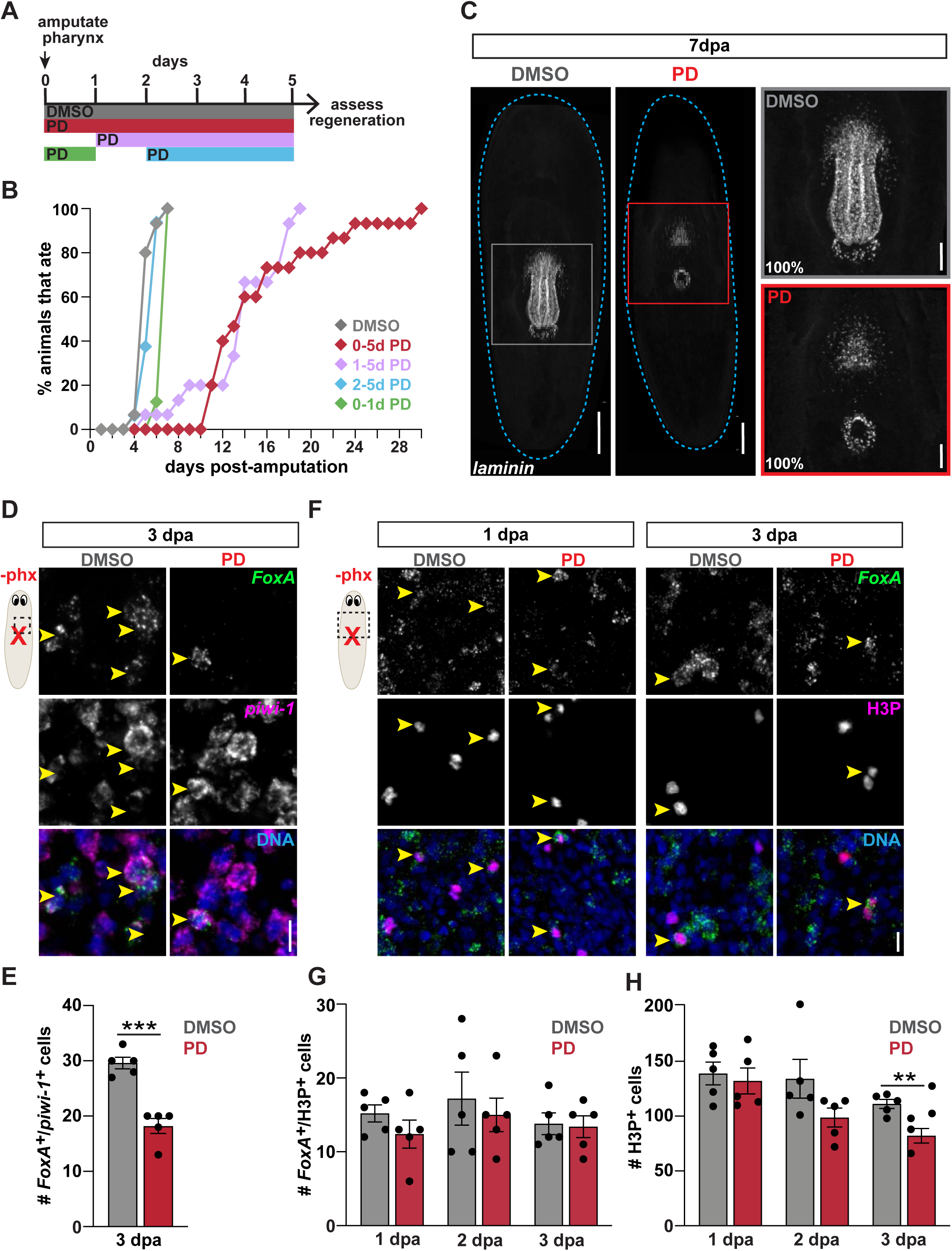
ERK Phosphorylation is Required for Increased Pharyngeal Progenitors. (A) Schematic of PD0325901 (PD) exposure relative to pharynx amputation for graph in B. (B) Graph of feeding behavior of animals after pharynx amputation, treated as indicated in A and assayed daily, represented as a proportion. n ≥ 20 animals. (C) Whole-mount FISH for the pharynx marker *laminin* 7 days after pharynx amputation in animals treated with DMSO or PD for 5 days. Dashed line outlines animal; boxes indicate zoomed area on right; scale bars = 250µm (left) or 50µm (right). n ≥ 10 animals. (D) Confocal Images of FISH for *FoxA* (green) and *piwi-1* (magenta) 3 days after pharynx amputation in animals treated with DMSO or PD. DNA = DAPI (blue); dashed boxes = region imaged; arrows = double-positive cells; scale bar = 10µm. (E) Graph of *FoxA*^*+*^ *piwi-1*^*+*^ cells in the area outlined by dashed boxes in D. (F) Confocal images of *FoxA* FISH (green) and H3P antibody (magenta) 1 and 3 days after pharynx amputation in animals treated with DMSO or PD. DNA = DAPI (blue); dashed boxes = region imaged; arrows = double-positive cells; scale bar = 10μm. (G) Graph of *FoxA*^*+*^ H3P^+^ cells in the area outlined by dashed boxes in F at different times after pharynx amputation in animals treated with DMSO or PD. (H) Graph of H3P^+^ cells in the area outlined by dashed boxes in F at different times after pharynx amputation in animals treated with DMSO or PD. Bar graphs are represented as mean ± SEM. In bar graphs: dots = individual animals; and **, p ≤ 0.01; ***, p ≤ 0.001, unpaired t-test.

To pinpoint when ERK signaling is important for pharynx regeneration during this 5-day window, we exposed animals to PD for defined times after pharynx amputation (Figure 4A) and then used feeding assays to monitor pharynx regeneration. Animals treated with PD beginning 1 day after pharynx amputation (1-5 days) display a similar delay in the ability to feed as animals treated for 5 full days. However, animals treated for only the first day after amputation (0-1 days) or beginning 2 days after pharynx amputation (2-5 days) regained the ability to feed at rates similar to controls (Figure 4B). These results define a window 1-2 days after amputation in which activation of ERK signaling is important for pharynx regeneration. Coincidentally, this time frame immediately follows the proliferation of *FoxA*^*+*^ stem cells (Figure 2C), but occurs just prior to the increase in pharynx progenitors (Figure 1H) observed after pharynx loss. The requirement for ERK activity in this window suggests that ERK may be important for establishing pharynx progenitors, but not necessarily for the proliferation of *FoxA*^*+*^ stem cells during regeneration.

Because ERK signaling has known roles in stem cell proliferation and differentiation in planaria (Owlarn et al., 2017; Tasaki et al., 2011), we tested whether ERK inhibition would impact the expansion of pharynx progenitors during pharynx regeneration. To determine whether ERK is required for generating pharynx progenitors, we maintained animals in PD for 3 days following pharynx amputation and performed FISH for *FoxA* and *piwi-1*. Exposure to PD caused a significant decrease in pharynx progenitors as compared to controls (Figure 4D and 4E). To determine whether this decrease was due to reduced proliferation, we analyzed the number of *FoxA*^*+*^ H3P^+^ stem cells at different times after pharynx amputation. The number of proliferating *FoxA*^*+*^ stem cells after exposure to PD was comparable to controls (Figure 4F and 4G). However, we did observe a slight, but significant, decrease in the total number of H3P^+^ cells 3 days after pharynx amputation, consistent with what has been reported in regenerating heads (Figure 4H) (Owlarn et al., 2017). This result indicates that ERK activation is not required for the specific increase in proliferation of *FoxA*^*+*^ stem cells caused by pharynx loss. Instead, ERK contributes to increased *FoxA* expression in pharynx progenitors, suggesting that it is required for stem cell differentiation during pharynx regeneration.

Soon after amputation, ERK signaling is required for expression of several genes including *follistatin (fst)* (Owlarn et al., 2017), which accelerates regeneration by inhibiting activin-1 and −2 (Gaviño et al., 2013; Roberts-Galbraith & Newmark, 2013; Tewari et al., 2018). Consequently, we predicted that ERK signaling could promote pharynx regeneration through induction of *fst* expression. After head amputation, *fst* expression increases within 6 hours (Gaviño et al., 2013). By contrast, we did not observe *fst* expression until 24 hours after pharynx amputation (Figure S3A), suggesting that pharynx removal may activate ERK later than head removal.

However, *fst*(*RNAi*) animals regained the ability to feed at a normal rate after pharynx amputation (Figure S3B), indicating that pharynx regeneration does not depend on *fst*. Therefore, although pharynx loss eventually induces *fst* expression, regulation of pharynx regeneration via ERK is independent of *fst*.

### Pharynx Loss Does Not Increase Non-pharyngeal Progenitors

Planarian progenitor stem cells can be distinguished by expression of various organ-specific transcription factors that are often required for subsequent organ regeneration. These include *ovo, myoD, gata4/5/6, six-1/2* and *pax6a*, which are also expressed in mature organs (eye, muscle, gut, excretory system and nervous system, respectively) (Figure 5A) (Flores et al., 2016; Lapan & Reddien, 2012; Rouhana et al., 2013; Scimone et al., 2011, 2017; Scimone, Kravarik, et al., 2014). Expression of these markers in *piwi-1*^+^ stem cells provides another opportunity to link the behavior of organ-specific progenitors with injury. Our data so far support a model in which pharynx tissue loss specifically amplifies *FoxA*^*+*^ pharynx progenitors. If this is true, head amputation, which disrupts all organ systems except the pharynx, should amplify non-pharyngeal progenitors while removal of the pharynx should not (Figure 5B). Conversely, a previous study showed that surgical amputation of the pharynx resulted in increased production of differentiated non-pharyngeal tissues, supporting a model in which injury non-specifically amplifies any nearby progenitors (Figure 5B) (LoCascio et al., 2017). However, the surgical removal of the pharynx employed in this study also injured surrounding tissues, making it difficult to decipher which injuries contributed to the production of differentiated cells. Therefore, to test these two models, we evaluated the behavior of multiple organ-specific progenitors after either pharynx or head amputation.

**Figure 5:**
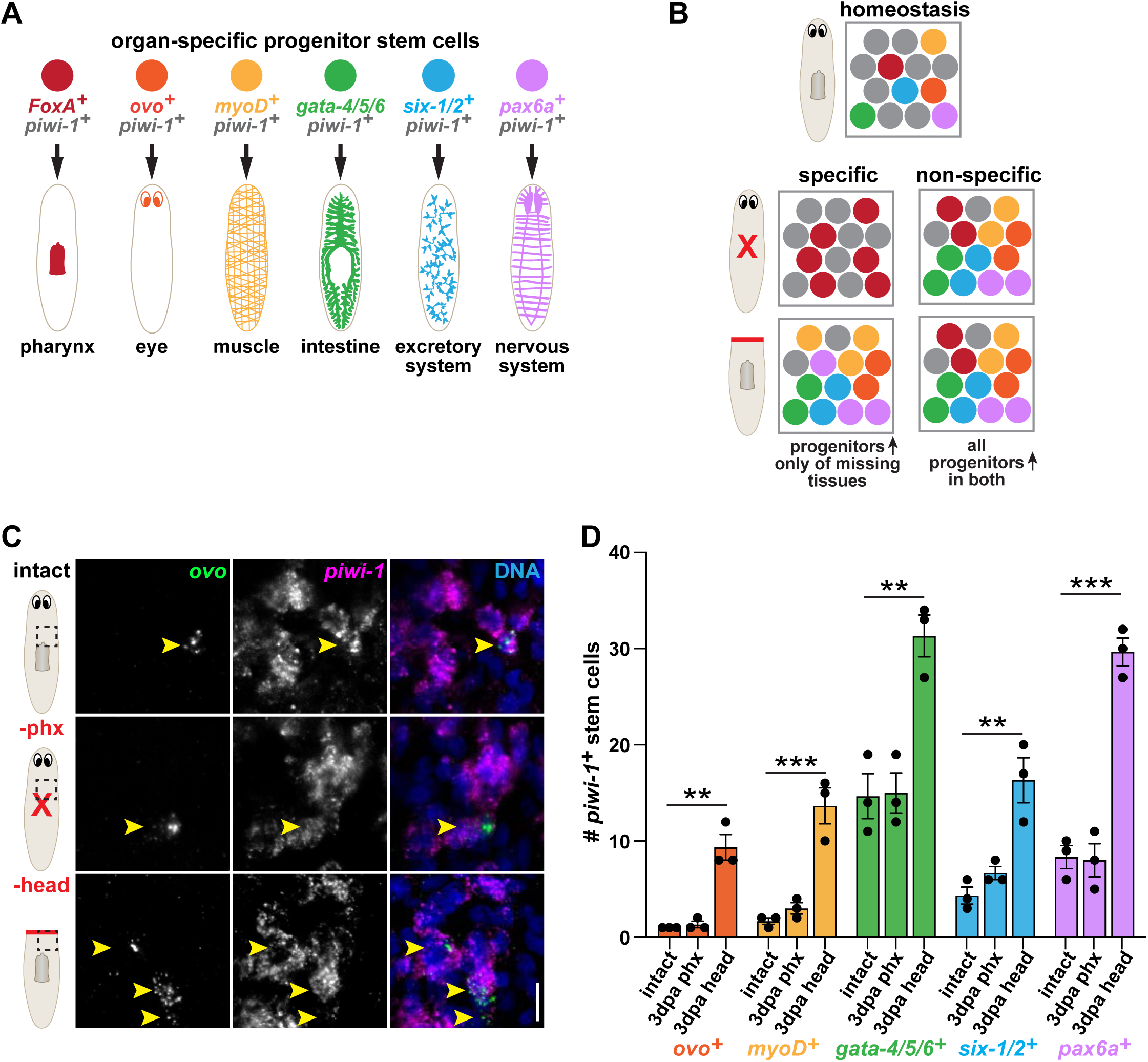
Pharynx Loss Does Not Increase Non-pharyngeal Progenitors. (A) Schematic of organ-specific progenitors and the organ systems they replenish. Progenitors express both *piwi-1* and organ-specific transcription factors. (B) Models for specific and non-specific organ regeneration. Organ-specific progenitor stem cells (colored as indicated in A), other stem cells (grey). (C) Confocal images of FISH for *ovo* (green) and *piwi-1* (magenta) in intact animals, or 3 days after pharynx or head amputation. DNA = DAPI (blue); dashed boxes = region imaged; arrows = double-positive cells; scale bar = 10µm. (D) Graph of cells double-positive for *piwi-1* and the indicated progenitor marker in the area outlined by dashed boxes in C, represented as mean ± SEM. Dots = individual animals; and **, p ≤ 0.01; ***, p ≤ 0.001, unpaired t-test.

Besides the pharynx, the eye is the only other organ in the planarian that is anatomically restricted and thus can be fully removed without leaving any remaining tissue behind (Lapan & Reddien, 2012; LoCascio et al., 2017). To test whether selective pharynx removal induces non-specific production of other tissue types, we first measured eye progenitor abundance 3 days after pharynx or head amputations with FISH for *ovo* and *piwi-1*. As previously shown, decapitated animals, where the eyes are completely removed, have significantly more eye progenitors than intact controls (Lapan & Reddien, 2012). Conversely, following pharynx amputation, animals had similar numbers of eye progenitors as intact controls (Figure 5C and 5D), indicating that pharynx loss does not stimulate the production of eye progenitors. If the broad damage caused by head amputation stimulates the specific regeneration of missing tissues, we predicted that an increase in other organ-specific progenitors after head, but not pharynx, amputation would also occur. To test this, we quantified organ-specific progenitors for muscle (*myoD*^*+*^), intestine (*gata-4/5/6*^*+*^), the excretory system (*six-1/2*^*+*^) and the nervous system (*pax6a*^*+*^) (Figure 5A) after either pharynx or head removal. These organ-specific progenitors increased 3 days after head removal, but not pharynx removal, as compared to intact controls (Figure 5D). These data indicate that pharynx loss does not stimulate production of non-pharyngeal progenitors. Also, our data suggests that organ-specific progenitors are induced to expand by removal of tissue from the organs that they replenish.

Pharynx, but not head removal, stimulates the proliferation of *FoxA*^*+*^ stem cells (Figure 2C and 2D). Stem cell progenitors in the epidermal lineage have also been shown to proliferate following head amputation (van Wolfswinkel et al., 2014). Therefore, we tested whether stem cells expressing non-pharyngeal progenitor markers were similarly stimulated to proliferate 1 and 2 days after pharynx or head amputation. We were unable to detect proliferating eye progenitors (*ovo*^*+*^ H3P^+^), even after head amputation (data not shown). However, stem cells expressing other progenitor markers did proliferate. The kinetics of each differed slightly, with increased proliferation of *gata-4/5/6*^*+*^ stem cells 1 day, and others (*six1/2*^*+*^, *pax6a*^*+*^, *myoD*^*+*^) 2 days after head amputation, but not pharynx amputation (Figure S4A and S4B). Despite slight differences in the kinetics of each progenitor marker, the overall trend supports the notion that only loss of non-pharyngeal tissues triggers proliferation of stem cells expressing non-pharyngeal progenitor markers.

All of these organ-specific transcription factors, with the exception of *pax6a*, are required for regeneration of their cognate organ (Adler & Sánchez Alvarado, 2017; Flores et al., 2016; Lapan & Reddien, 2012; Pineda et al., 2002; Scimone et al., 2011, 2017). To verify that these transcription factors do not regulate pharynx regeneration, we knocked them down with RNAi. However, knockdown did not impact the recovery of feeding behavior after pharynx amputation (Figure S4C). Together, these data suggest that tissue loss specifically amplifies only the organ-specific progenitors that are required to regenerate missing tissues.

### Proliferation and ERK Activation are Not Required for Eye Regeneration

Decapitated head fragments lack a pharynx, but retain eyes, providing a different context in which to investigate the specificity of stem cell responses to missing organs. Therefore, we analyzed pharynx and eye progenitor behavior in decapitated head fragments. As expected, the number of pharynx progenitors in head fragments was significantly higher 3 days after amputation. Conversely, eye progenitors showed no discernible difference between 0 and 3 days after amputation (Figure 6A and 6B). This lack of eye progenitor amplification in head fragments was somewhat surprising because previous studies have shown that tissue removal near the eye, without removing the eye itself, increases eye progenitors (LoCascio et al., 2017). Regardless, these results further support a model in which stem cells modify their behavior depending on what organs are present or absent.

**Figure 6:**
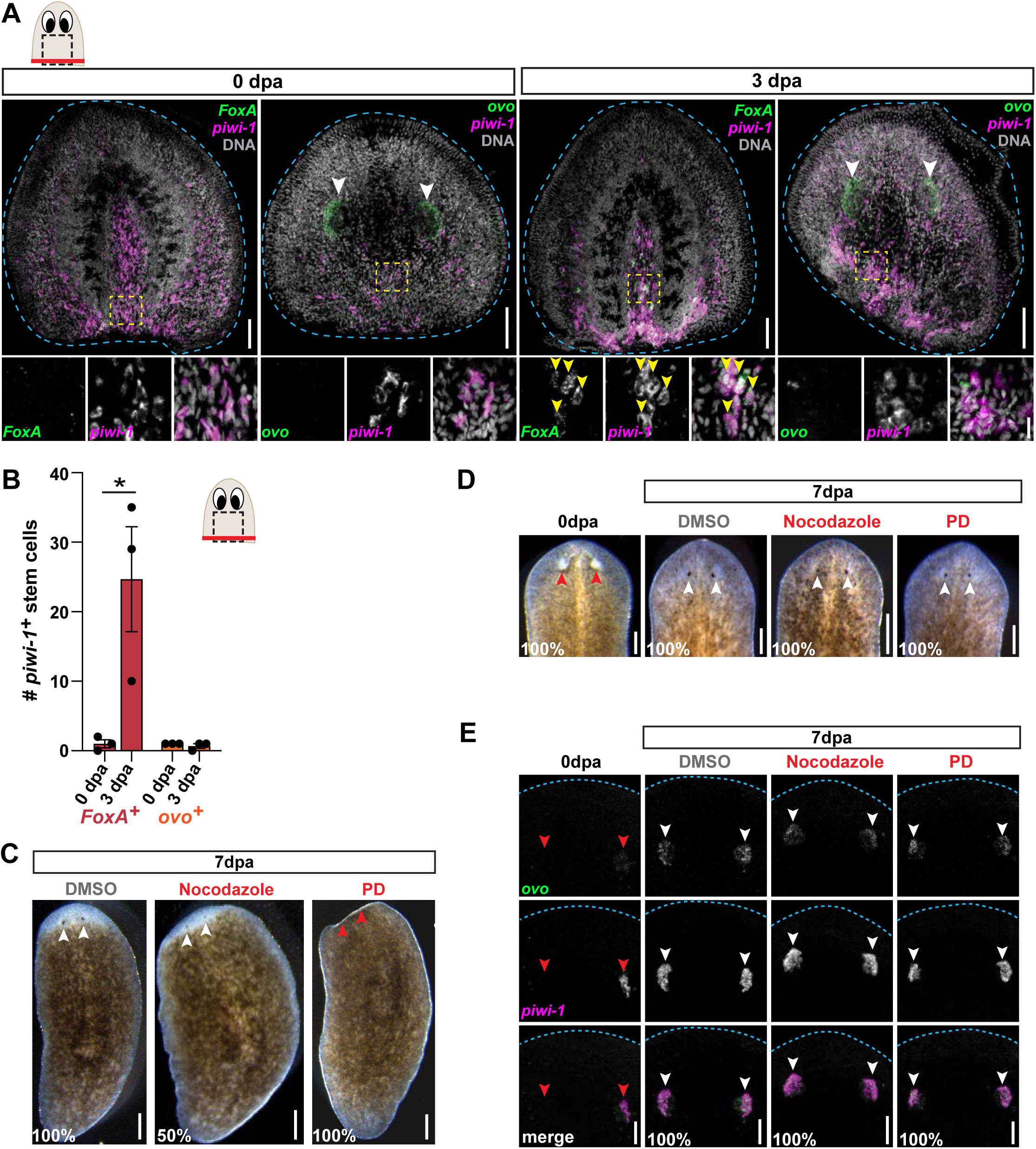
Proliferation and ERK Activation are Not Required for Eye Regeneration. (A) Confocal images of FISH for *piwi-1* (magenta) and *FoxA* or *ovo* (green) in head fragments 0 and 3 days after amputation. Blue dashed line outlines head fragment; DNA = DAPI (grey); dashed boxes = zoomed area below; white arrows = eyes; yellow arrows = double-positive cells; scale bars= 50µm (top) and 10µm (top). (B) Graph of *FoxA*^*+*^ *piwi-1*^*+*^ and *ovo*^*+*^ *piwi-1*^*+*^ cells in the area outlined by dashed boxes in the cartoon, represented as mean ± SEM. Dots = individual animals; and *, p ≤ 0.05, unpaired t-test. (C) Live images 7 days after head amputation of animals treated with PD or DMSO for 5 days after amputation, or nocodazole for 1 day beginning 24 hours after amputation. Red arrows = no eyes, white arrows = regenerated eyes; scale bars = 250µm. n ≥ 10 animals. (D) Live images 7 days after eye resection in animals treated with PD or DMSO for 5 days, or nocodazole for 48 hours, immediately after resection. Red arrows = resected eyes; white arrows = regenerated eyes; scale bars = 50µm. n ≥ 10 animals. (E) Confocal images of FISH for *ovo* (green) and *opsin* (magenta) 7 days after eye resection in animals treated with PD or DMSO for 5 days, or nocodazole for 48 hours immediately after resection. Red arrows = resected eyes; white arrows = regenerated eyes; scale bars = 50µm. n ≥ 10 animals.

Selective removal of the eye does not increase broad stem cell proliferation or eye-specific progenitors (LoCascio et al., 2017), and we were unable to detect proliferating *ovo*^*+*^ stem cells even after head amputation (data not shown). To test whether eye regeneration relies on proliferation, as pharynx regeneration does, we exposed worms to nocodazole for 24 hours beginning 1 day after head amputation, and assessed the emergence of photoreceptors. In DMSO-treated controls, photoreceptors re-emerged 7 days after amputation (Figure 6C) (Lapan & Reddien, 2011). While nocodazole-treated worms failed to regenerate heads and had noticeably smaller blastemas than control animals, photoreceptors still emerged by 7 days in 50% of animals. By contrast, consistent with what has been published, animals treated with the ERK inhibitor, PD, for 5 days after head removal completely failed to regenerate any new head tissue (Figure 6C) (Owlarn et al., 2017; Tasaki et al., 2011). These data suggest that eyes can regenerate after decapitation without stem cell proliferation.

To further test the requirement for proliferation and ERK signaling during eye regeneration, we evaluated eye regeneration after more subtle surgeries in which the eyes are selectively removed. Following eye resections, we immediately exposed animals either to nocodazole or PD and monitored eye regeneration with live imaging and FISH for *ovo* and the eye-specific marker *opsin* (A. Sánchez Alvarado & Newmark, 1999). We confirmed that eye tissue was successfully removed by the absence of photoreceptors in animals 0 days after surgery (Figure 6D). While FISH revealed that *opsin* staining was not completely removed, it was strongly diminished and *ovo* signal was eliminated (Figure 6E). Even when worms were exposed to nocodazole for 48 hours, which caused some animal lethality, eyes regenerated normally in surviving worms. Animals treated with PD for 5 days following resection also regenerated eyes similar to DMSO-treated controls (Figure 6D and 6E). Therefore, unlike pharynx regeneration, eye regeneration does not require proliferation and only requires ERK if more tissue has been removed, such as in the context of head regeneration. These data suggest that eye regeneration may be regulated differently than other organs, or may occur in alternative ways depending on the context of the wound.

## Discussion

In this study, we challenged the planarian stem cell compartment by inflicting different types of injuries to evaluate how stem cells sense organ loss and initiate tissue-specific regeneration. We show that proliferation and expansion of organ-specific progenitors depends on the removal of tissues that they are required to regenerate (Figure 7). In particular, we find that pharynx regeneration depends on an increase in pharynx progenitors that occurs only if pharynx tissue is lost. This increase first relies on a burst in proliferation of *FoxA*^*+*^ stem cells, followed by ERK signaling that likely drives differentiation of stem cells into pharynx progenitors. Conversely, stem cells expressing non-pharyngeal progenitor markers divide and increase only when the tissues they are required to regenerate are removed. While head regeneration is also dependent on proliferation and ERK activation, regeneration of the eyes is not (Figure 7). These findings suggest that in many cases, stem cells can sense the identity of missing tissues to launch their targeted regeneration.

**Figure 7:**
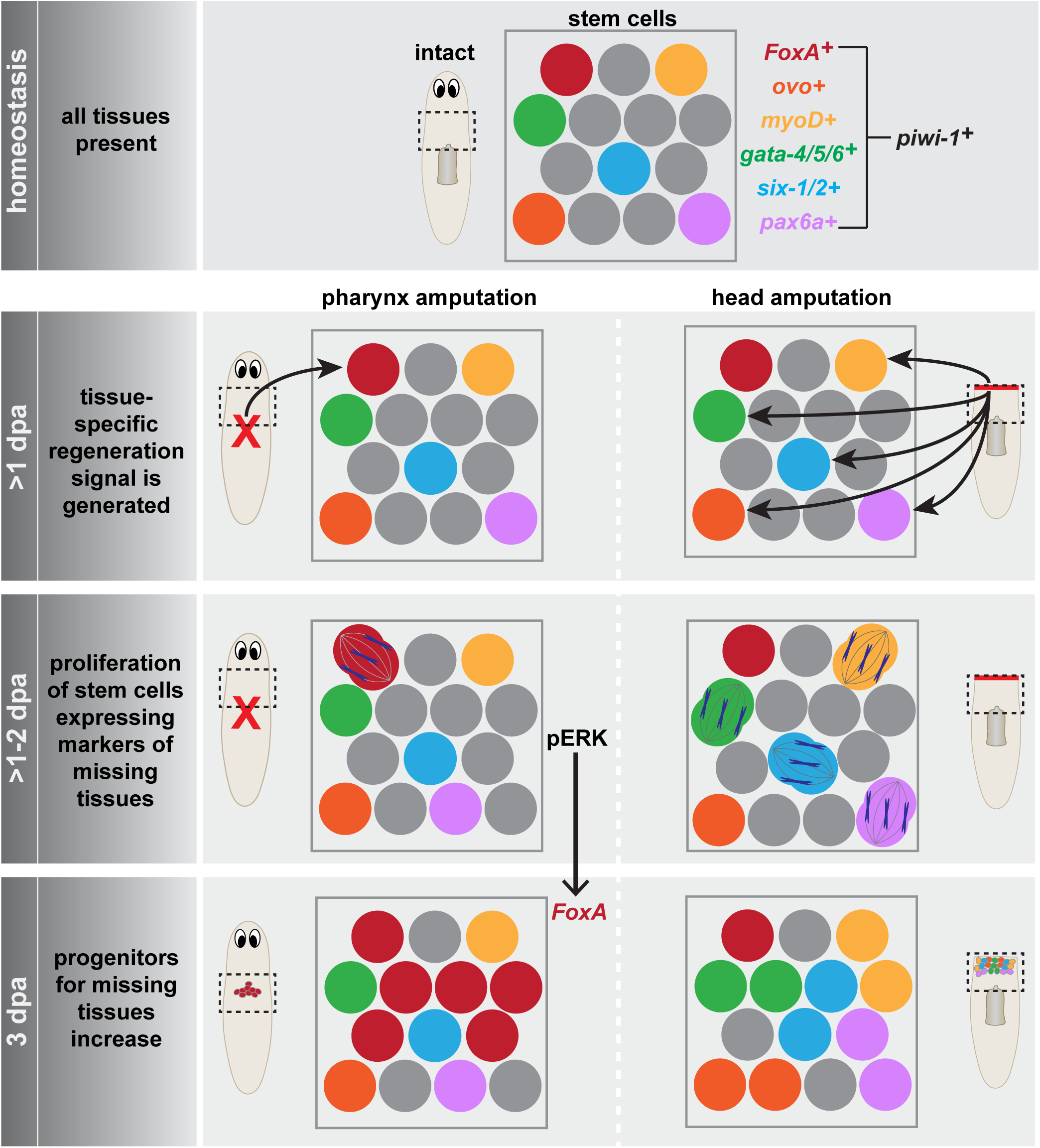
Model for Amplification of Progenitors During Regeneration. In intact animals during homeostasis, organ-specific progenitor markers are expressed in a subset of stem cells. Within 1 day of either pharynx or head amputation, stem cells expressing progenitor markers for missing tissues (except *ovo*) are induced to proliferate. Three days after amputation, progenitors specific to missing tissues increase in number. *FoxA*^+^ stem cells proliferate and subsequently increase pharynx progenitors, dependent on ERK signaling.

### Stem Cell Heterogeneity and Specificity of Regeneration

Molecular heterogeneity is a feature of many stem cell systems including embryonic, neural, and muscle stem cells (Chaker et al., 2016; Scaramozza et al., 2019; Torres-Padilla & Chambers, 2014). While planarian PSCs are sufficient to reconstitute entire animals after transplantation by producing the necessary organ-specific progenitors (Wagner et al., 2011; Zeng et al., 2018), how stem cell heterogeneity is utilized during regeneration has been difficult to decipher. Traditional amputation methods injure multiple tissues at once, are typically variable in position and size, and also cause major disruptions to positional patterning of the anterior-posterior axis, complicating the study of regeneration (Reddien, 2018). Selective removal of the pharynx circumvents these issues in two ways. First, the tissue removed by chemical amputation and the wound that it generates are reproducible among animals. Second, it does not perturb axial patterning or other organ systems. Together, these advantages enable us to challenge the stem cell population in a precise way. By studying the dynamics of *FoxA* expression in stem cells after pharynx removal, we have uncovered shifts in heterogeneity that depend on the presence or absence of a particular organ. Intriguingly, we find that stem cells expressing both the PSC marker, *tgs-1*, and *FoxA*, or *FoxA* alone are stimulated to divide after pharynx removal. Therefore, both PSCs and pharynx progenitors respond to pharynx loss and channel their proliferative output towards pharynx regeneration. Combined with our analysis of non-pharyngeal progenitor dynamics after different amputations, our results suggest that the heterogeneity of the stem cell population can be differentially deployed depending on what tissues need repair.

Previous work showed that large wounds stimulate broad differentiation of stem cells into nearby organs, while eye resection alone does not stimulate an increase in *ovo*^*+*^ eye progenitors (LoCascio et al., 2017). Based on these observations, the authors speculated that stem cells non-specifically modify their output of progenitors based on the size and position of a wound, instead of the identity of missing tissues. However, by investigating the dynamics of a wide variety of organ-specific progenitors after different injuries, we show that stem cells can adopt specific, tailored responses to the loss of different tissues. Amputation of non-pharyngeal tissues increases non-pharyngeal progenitors, while pharynx tissue loss triggers a specific increase of pharynx progenitors. Therefore, although some aspects of regeneration are likely to be general, amplification of progenitors typically requires the removal of the tissues they are required to regenerate. Smaller lesions, such as eye resections, do not stimulate a proliferative wound response (LoCascio et al., 2017). This difference could potentially explain why eye regeneration, in this context, happens passively via homeostatic progenitor production and reduced apoptosis in the regenerating eye (LoCascio et al., 2017). It is likely that stem cells increase proliferation rates, and differentially upregulate organ-specific transcription factors, depending on the type of damage inflicted to the animal. This dynamic response of stem cells may enable a flexible cellular output to replenish damaged tissues.

### Contribution of Proliferation to Regeneration

In many organisms, elevated proliferation at sites of injury is a prominent feature of regeneration (Tanaka & Reddien, 2011). In planaria, an initial body-wide wave of proliferation occurs approximately 6 hours after any type of injury, followed by a second local proliferative wave at 48 hours, but only if tissue has been removed (Baguñà, 1976; Wenemoser & Reddien, 2010). Little is known about how and when particular cell types are produced during each of these waves of proliferation. By pairing pharynx removal with *FoxA* expression in stem cells, we have dissected the contribution of proliferation to subsequent organ regeneration. Our observation that an increase in *FoxA*^*+*^ proliferating stem cells appears robustly within 6 hours of pharynx, but not head amputation, indicates that stem cells sense the identity of missing tissues during this first wave of proliferation. Because injury is only sufficient to induce regeneration when tissue is missing (Owlarn et al., 2017), all injuries may generate the same signals but only trigger regeneration under certain circumstances. Consistently, our findings suggest that missing tissue signals cause selective amplification of pharynx progenitors to confer a specific regenerative outcome when the pharynx is lost. Additionally, loss of non-pharyngeal tissues stimulates proliferation of stem cells expressing non-pharyngeal progenitor markers. These data suggest that missing tissue signals may be a general strategy used to channel the fate of stem cells to replace damaged organs during initial proliferative responses.

By preventing mitotic exit with nocodazole, we made the surprising discovery that proliferation in a narrow window of time following amputation is absolutely necessary for pharynx regeneration. Nocodazole exposure for 24 hours beginning one day after amputation severely delayed the recovery of feeding behavior. Surprisingly, nocodazole exposure for a 24-hour period either before or after this window did not have a similar effect, indicating that cell division 2-3 days after amputation is not required for pharynx regeneration. This time frame is concurrent with the peak of proliferation that occurs after tissue removal, and is consistent with a recent finding that the missing tissue response is not required for regeneration (Tewari et al., 2018). Further, nocodazole exposure completely blocked the increase of pharynx progenitors that typically occurs 3 days after pharynx amputation. Therefore, without proliferation in this narrow window, cell production required to rebuild the pharynx occurs at a significantly slower rate. Because regeneration depends on proliferation during such a short time, our results suggest that stem cells detect tissue loss through transient signals generated by injury that are necessary to accelerate subsequent regeneration.

How or when fate acquisition might occur in these proliferating cells during regeneration remains unclear. Cell fate acquisition can occur throughout the cell cycle (Fichelson et al., 2005; Pauklin & Vallier, 2014; Soufi & Dalton, 2016). Tissue loss could generate fleeting signals sensed by stem cells that influence them to adopt a specific cell fate during proliferation to compensate for missing tissue. Alternatively, progenitors may be poised to receive such a signal, allowing them to quickly initiate regeneration upon exit of the cell cycle. Indeed, studies in human hepatoma cell lines have shown that FoxA1 remains attached to chromatin during mitosis, contributing to rapid activation of downstream following mitosis during liver differentiation (Caravaca et al., 2013). Our data show that both *FoxA*^*+*^ pharynx progenitors and PSCs expressing *FoxA* increase proliferation after pharynx removal, suggesting that a combination of both might be true. Future work will define whether regeneration of other organs also depends on proliferation in such a narrow window, as well as the nature of the signals responsible for selectively amplifying different progenitor types.

### ERK Signaling Plays Multiple Roles During Regeneration

Phosphorylation of ERK activates regeneration in many animals (DuBuc et al., 2014; Wan et al., 2012; Yun et al., 2014). In planaria, ERK phosphorylation is one of the earliest known events required for regeneration, where it triggers wound-induced transcription and promotes stem cell proliferation and differentiation (Owlarn et al., 2017; Tasaki et al., 2011). ERK may also function to re-establish axial patterning during regeneration (Owlarn et al., 2017; Umesono et al., 2013), which depends on a network of positional cues that are expressed in muscle cells throughout the body (Lander & Petersen, 2016; Scimone et al., 2016; Witchley et al., 2013). Because injuries that disrupt muscle require re-establishment of these positional cues for regeneration to proceed (Rink, 2018), it has been difficult to distinguish ERK’s functions in initiating organ regeneration with its roles in axial re-patterning in studies where bodily injuries were employed. Removal of the pharynx does not disrupt body wall muscle, which allowed us to identify a distinct role for ERK in organ regeneration. Following pharynx removal, ERK activation is not involved in proliferation of *FoxA*^*+*^ stem cells but is required for the subsequent increase in pharynx progenitors. These results show that another signal must trigger proliferation of *FoxA*^*+*^ stem cells, and that ERK acts later during organ regeneration, likely to facilitate stem cell differentiation or to maintain progenitor fate.

Receptor tyrosine kinases such as the epidermal growth factor receptor (EGFR) and the fibroblast growth factor receptor (FGFR) have been shown to play critical roles in signaling upstream of ERK in many organisms (Patel & Shvartsman, 2018), making them intriguing candidates to explore in planaria as potential regulators of regeneration. In planarians, *egfr-3* is required to activate ERK during regeneration (Fraguas et al., 2017) and is also involved in stem cell differentiation (Fraguas et al., 2011; Lei et al., 2016). Other studies have also highlighted roles for the ligand *egf-4* and the receptors, *egfr-1* and *egfr-5*, in the differentiation of stem cells into brain, intestinal and excretory tissues, respectively (Barberán et al., 2016; Fraguas et al., 2014; Rink et al., 2011). Whether any of the planarian EGF or FGF ligands or receptors similarly regulate the differentiation of pharyngeal progenitors is an area for future investigation (Cebrià et al., 2002; Ogawa et al., 2002).

### Pioneer Factors: Manifesting Specific Regeneration Programs in Stem Cells

Following pharynx loss, planarian *FoxA* is selectively upregulated in stem cells to drive organ regeneration. Mammalian homologs of *FoxA* were the first identified ‘pioneer’ transcription factors, characterized by their ability to engage closed chromatin and drive organogenesis (Hsu et al., 2015; Iwafuchi-Doi & Zaret, 2016; Lam et al., 2013; Zaret & Mango, 2016). This raises the possibility that pioneer factors may be viable *in vivo* targets for achieving regeneration of entire organs. In fact, overexpression of a related mammalian transcription factor, *FoxN*, is sufficient to drive regeneration of the thymus in mice (Bredenkamp et al., 2014). The increased proliferation of PSCs expressing *FoxA* after pharynx removal suggests that activation of pioneer factors in stem cells may also drive organ regeneration in planaria. Other pioneer factors, including *gata-4/5/6, soxB1-2* and *FoxD*, are also expressed in planarian stem cells and are required for regeneration of the intestine (Flores et al., 2016; González-Sastre et al., 2017), sensory neurons (Ross et al., 2018) and anterior pole (Scimone, Lapan, et al., 2014; Vogg et al., 2014), respectively. Therefore, upregulation of pioneer factors in stem cells may be a general strategy used to initiate organ regeneration. Identifying the regulatory mechanisms responsible for the selective activation of pioneer factors in stem cells may be an ideal approach to understanding how organisms initiate regeneration of targeted organs *in vivo*.

## Acknowledgements

We thank the Cornell University Biotechnology Resource Center for assistance with data collection on the Zeiss LSM 710 Confocal which is supported by the NIH (NIH S10RR025502). We would also like to thank Kuang-Tse Wang for helpful comments on the manuscript. D.A.S. was supported by a GRA Fellowship from the Cornell College of Veterinary Medicine. This work was supported by C.E.A.’s Cornell University startup funds.

## Author Contributions

Conceptualization, T.E.B., D.A.S., C.E.A.; Methodology, T.E.B. and D.A.S.; Investigation, T.E.B.; Writing – Original Draft, T.E.B. and C.E.A.; Writing – Review & Editing, T.E.B., D.A.S., C.E.A.; Supervision, C.E.A., Funding Acquisition, C.E.A.

## Lead Contact

Further information and requests for resources and reagents should be directed to and will be fulfilled by the Lead Contact, Carolyn Adler.

## Materials Availability

Plasmids generated in this study will be made available on request, without restriction.

## Data and Code Availability

This study did not generate/analyze [datasets/code]

## Supplemental Figure Legends

Supplementary Figure 1: ***FoxA***^***+***^ **Pluripotent Stem Cells Proliferate after Pharynx Amputation**

(A) Confocal images of *FoxA* (turquoise) and *tgs-1* (magenta) FISH, and H3P antibody (yellow) in intact animals, and 1 day after pharynx amputation. DNA = DAPI (grey); yellow arrows = only H3P+; blue arrows = *FoxA* H3P double-positive; white arrows = *FoxA, tgs-1* and H3P triple-positive; scale bar = 10μm.

(B) Graph of H3P^+^ cells that are *FoxA*^*-*^ *tgs-1*^*+*^ (green), *FoxA*^*+*^ *tgs-1*^*-*^ (red), and *FoxA*^*+*^ *tgs-1*^*+*^ (yellow) 1 day after pharynx amputation in the area outlined by dashed boxes in A, represented as mean ± SEM. Dots = individual animals; and *, p ≤ 0.05; ***, p ≤ 0.001, unpaired t-test.

Supplementary Figure 2: **Nocodazole Treatment Stalls *FoxA***^***+***^ **Stem Cells in Mitosis after Pharynx Amputation**

(A) Confocal images of *FoxA* FISH (green) and H3P antibody (magenta) 2 days after pharynx amputation in animals treated with DMSO or nocodazole, 1 day after amputation for 24 hours. DNA = DAPI (blue); arrows = double-positive cells; scale bar = 10μm.

(B) Graph of *FoxA*^*+*^ H3P^*+*^ cells quantified from A, represented as mean ± SEM.

(C) Confocal images of FISH for *FoxA* (green) and *piwi-1* (magenta) in intact animals treated with the DMSO or nocodazole, for 24 hours and fixed 1 day after wash out. DNA = DAPI (blue); arrows = double-positive cells; scale bar = 10μm.

(D) Graph of *FoxA*^*+*^ *piwi-1*^*+*^ cells from C. Graphs represented as mean ± SEM. Dots = individual animals; and ***, p ≤ 0.001, unpaired t-test.

Supplementary Figure 3: **ERK-dependent Pharynx Regeneration is Independent of *follistatin***

(A) Whole-mount colorimetric *in situ* hybridization of *fst* in intact animals and at different times after pharynx or head amputation. Arrows = amputation site; scale bars = 250µm.

(B) Feeding behavior in control (*unc-22*), *FoxA* and *fst* RNAi animals 7 days after pharynx amputation, represented as a proportion of the total. n ≥ 10 animals.

Supplementary Figure 4: **Proliferation of Stem Cells Expressing Non-pharyngeal Progenitor Markers is Not Triggered by Pharynx Loss**

(A) Graph of cells double-positive for H3P and the indicated organ-specific progenitor marker in intact animals, or 1 day after pharynx or head amputation. Dashed boxes = region quantified.

(B) Graph of cells double-positive for H3P and the indicated organ-specific progenitor marker in intact animals, or 2 days after pharynx or head amputation. Dashed boxes = region quantified.

(C) Graph of feeding behavior 7 days after pharynx amputation in RNAi animals, represented as a proportion of the total. n ≥ 10 animals. Bar graphs represented as mean ± SEM. Dots = individual animals; and *, p ≤ 0.05; **, p ≤ 0.01, unpaired t-test.

## Materials and Methods

### Worm care

*Schmidtea mediterranea* asexual clonal line CIW4 were maintained in a recirculating system which facilitates constant UV sanitization of water containing Montjuïc salts (planaria water) (Arnold et al., 2016; Merryman et al., 2018). Prior to their use for experiments, animals were transferred to static culture and maintained in planaria water supplemented with 50 µg/mL gentamicin sulfate. Animals used for experiments were between 2-3mm in length and starved for approximately 5-7 days.

### Amputations, sodium azide treatment and tricaine anesthetization

Amputations were done on 2-3 mm animals. Selectively pharynx removal was performed by chemical amputation as previously described (Adler et al., 2014; Shiroor et al., 2018). Planarians were placed in 100mM sodium azide diluted in planaria water. After 4-7 min, the pharynx extended out of the body and was plucked off using fine forceps (#72700-D; Electron Microscopy Sciences). The sodium azide was removed no more than 10 min after initial exposure and worms were rinsed 3 times, removing all loose pharynges, before being transferred into a fresh dish. Heads were removed using a micro feather scalpel (#72045-45; Electron Microscopy Sciences) to remove the head from the body approximately halfway between the eyes and the pharynx. When compared to chemical pharynx amputations, head amputations were performed and intact animals were soaked in sodium azide for 2-3 minutes. For pharynx incisions and partial amputations, animals were soaked in tricaine solution (4g/L in 21mM Tris pH 7.5) diluted 1:3 in planaria water which causes the pharynx to extrude but not eject it. Pharynx incisions were created by inserting forceps into the pharynx and snipping along its length. To remove a portion of the pharynx, the proximal end of the pharynx was stabilized with forceps while a scalpel trimmed off the distal end. Eye resections were carried out by immobilizing animals on moist filter paper on a cooled block. The tips of fine forceps were used to scrape out the eye.

### *In situ* hybridizations, immunostaining and microscopy

Animals were fixed as previously described (Pearson et al., 2009) with minor modifications. Briefly, animals were killed in 7.5% N-acetyl-cysteine in PBS for 7.5 minutes and fixed in 4% paraformaldehyde in PBSTx (PBS + 0.3% Triton X-100) for 30 minutes. Worms were then rinsed twice with PBSTx and incubated in pre-warmed reduction solution (PBS+ 1% NP-40 + 50mM DTT + 0.5% SDS) at 37°C for 10 minutes. Worms were rinsed twice more with PBSTx, dehydrated in a methanol series and stored at −20°C.

Colorimetric *in situ* hybridizations were performed as described in (Pearson et al., 2009) using the anti-DIG-AP (Roche 11093274910) at 1:3000. Fluorescent *in situ* hybridizations were performed as in (King & Newmark, 2013) with minor modifications. Briefly, probes were generated for the genes indicated below with the provided animals were rehydrated and bleached (5% formamide, 1.2% H_2_O_2_ in 0.5x SSC) for 2 hours, then treated with proteinase K (4 µg/mL in PBSTx, Thermo Fisher 25530049). Following overnight hybridizations at 56°C, samples were washed 2x each in wash hybe (5 min), 1:1 wash hyb:2X SSC-0.1% Tween 20 (10 min), and 2X SSC-0.1% Tween 20 (30 min), 0.2X SSC-0.1% Tween 20 (30 min) at 56°C followed by 3 x 10 minute PBSTx washes at room temperature. Subsequently, animals were placed in blocking solution (0.5% Roche Western Blocking Reagent and 5% inactivated horse serum for POD and AP antibodies and 0.2% BSA for DNP antibodies diluted in PBSTx). Animals were then incubated with an appropriate antibody: 1:1000 anti-DIG-POD (Roche 11207733910), 1:1000 anti-FITC-POD (Roche 11426346910) or 1:200 anti-DNP-HRP (Perkin Elmer FP1129) in blocking solution at 4°C overnight (2 nights for anti-DNP). Antibodies were washed off in PBSTx and probes were developed by pre-incubation with tyramide (1:2000 FAM; 1:7,500 Cy3) in borate buffer for 30 minutes and then developed with 0.005% H_2_O_2_ in borate buffer for 45 minutes. Animals were pre-incubated with rhodamine tyramide (1:5000) for 10 minutes and developed for 15. For the development of subsequent probes, or detection of H3P, peroxidase was inactivated with 200mM sodium azide in PBSTx for 1 hour, then rinsed with PBSTx >6 times before application of the next antibody.

Animals were stained with phosphohistone H3 following *in situ* hybridizations by incubation in anti-phosphohistone H3 (Ser10) antibody (Abcam, Cambridge, MA Ab32107) at a concentration of 1:1000 in blocking solution (0.5% Roche Western Blocking Reagent and 5% inactivated horse serum in PBSTx) for 2 days at 4°C. Primary was washed off with PBSTx followed by incubation with either goat anti-rabbit-Alexa Fluor 647 (Life Technologies A21072) at 1:500 or goat anti-rabbit-HRP (Thermo Fisher 31460) at 1:2000 in PBSTx overnight at 4°C. Antibody was washed off with PBSTx and samples incubated in HRP antibody were pre-incubated in rhodamine tyramide (1:5000 in PBSTx) for 10 minutes and then developed with 0.005% H_2_O_2_ in PBSTx for 15 minutes.

DAPI [5µg/mL] (Thermo Scientific) diluted 1:5000, was added during the last antibody incubation. After development, animals were soaked in ScaleA2 (4M urea, 20% glycerol, 0.1% Triton X-100, 2.5% DABCO) (Hama et al., 2011) for at least 3 days and mounted in Aqua-Polymount (Polysciences Inc. 18606). Whole-mount colorimetric *in situ* hybridizations and live worms were imaged on a Leica M165F. Fluorescent *in situ* hybridizations were imaged on a Zeiss 710 confocal microscope. Images were processed in Fiji (Schindelin et al., 2012).

### Nocodazole treatment

Nocodazole (Sigma M1404) was administered in 24 or 48 hour increments at 50ng/mL diluted in planaria water containing 0.05% DMSO. Following treatment, animals were rinsed 3 times, transferred to a fresh dish and rinsed daily until further experimentation.

### PD0325901 MEK1/2 inhibitor III (PD) treatment

PD0325901 (EMD Millipore™ Calbiochem™ 4449685MG) was administered at 10µM diluted in planaria water containing 0.02% DMSO, and replaced daily. Animals whose treatment started at 0 days post-amputation were also soaked in PD 1 hour prior to and during amputation. After removal of PD, animals were rinsed 3 times and either fixed immediately, or transferred to a new dish and rinsed daily until further experimentation.

### Feeding assay

Animals were delivered 20µL of colored food (4:1 liver:milliQ water with 2% red food coloring) in a petri dish. Percentage of animals with red intestines were scored after approximately 30 min of food exposure. For time courses, feeding assays started at 4 days post-amputation and any animals that ate were scored and then removed from the dish. Feeding assay time courses were repeated at least three times with ∼20 animals assayed per experiment.

### Feeding RNAi

RNAi was carried out as previously described (Rouhana et al., 2013), with a few exceptions. Briefly, double stranded RNA (dsRNA) was synthesized *in vitro* using PCR products of mentioned genes. dsRNA was then diluted in colored food to a final concentration of 400ng/µL. Animals were removed from the incubator and immediately fed every 3 days, for a total of 6 feeds, except for *gata-4/5/6* and *six-1/2* RNAi which developed phenotypes after 1-2 feeds. *C. elegans unc22* dsRNA was used as a control. Amputations were carried out 5-7 days after the last feed. All RNAi experiments were repeated at least twice with at least 10 animals per experimental group. Efficient knockdown was confirmed by WISH and/or manifestation of known phenotypes.

### Quantification and statistical analysis

Quantification of *piwi-1*^*+*^ progenitors was performed with a minimum of 3 animals for each experimental group. Quantification was carried out manually in size-matched areas in the same approximate region of the animal. Quantification of H3P was performed in 5 animals for each experimental group. The anterior trunk region was imaged for H3P-stained animals and all H3P foci in the imaged region were quantified manually. In all bar graphs, dots represent individual animals and error bars represent standard error. Statistical analysis was performed using PRISM-Graphpad version 8 to perform an unpaired t-test. *p ≤ 0.05; **p ≤ 0.01, ***p ≤ 0.001 and ****p ≤ 0.0001.

### Gene annotation

Sequences for all transcripts used in this study for *in situ* hybridization and dsRNA synthesis were previously published and were identified in the ‘dd_Smed_v6’ transcriptome in PlanMine (Brandl et al., 2016) as follows: *piwi-1* (dd_Smed_v6_659_0_1) (Reddien et al., 2005); *FoxA* (dd_Smed_v6_10718_0_4) (Adler & Sánchez Alvarado, 2015; Scimone, Kravarik, et al., 2014); *tgs-1* (dd_Smed_v6_10988_0_1) (Zeng et al., 2018); *laminin* (dd_Smed_v6_8356_0_1) (Cebrià & Newmark, 2007); *fst* (dd_Smed_v6_9584_0_1) (Wenemoser et al., 2012); *ovo* (dd_Smed_v6_48430_0_1) (Lapan & Reddien, 2012); *myoD* (dd_Smed_v6_12634_0_1) (Cowles et al., 2013); *gata-4/5/6* (dd_Smed_v6_4075_0_1) (Wagner et al., 2011); *six-1/2* (dd_Smed_v6_9774_0_1) (Scimone et al., 2011); and *pax6a* (dd_Smed_v6_17726_0_1) (Pineda et al., 2002).

