## Supplementary figures and images for "Planarian stem cells sense the identity of missing tissues to launch targeted regeneration"

### Supplemental Figures 1-4

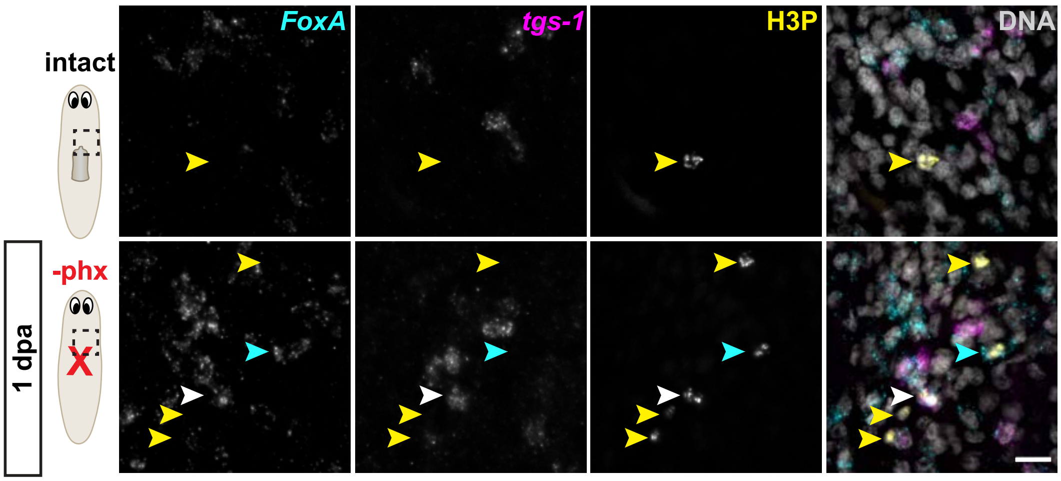

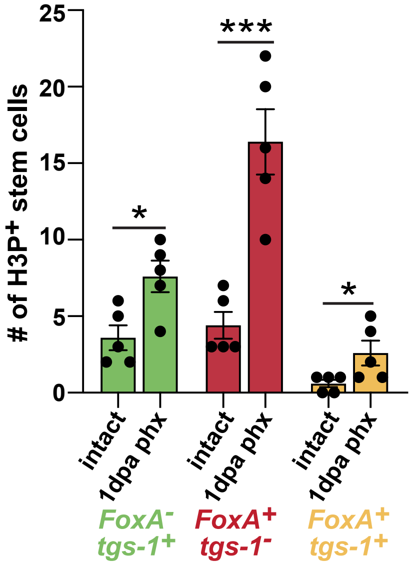

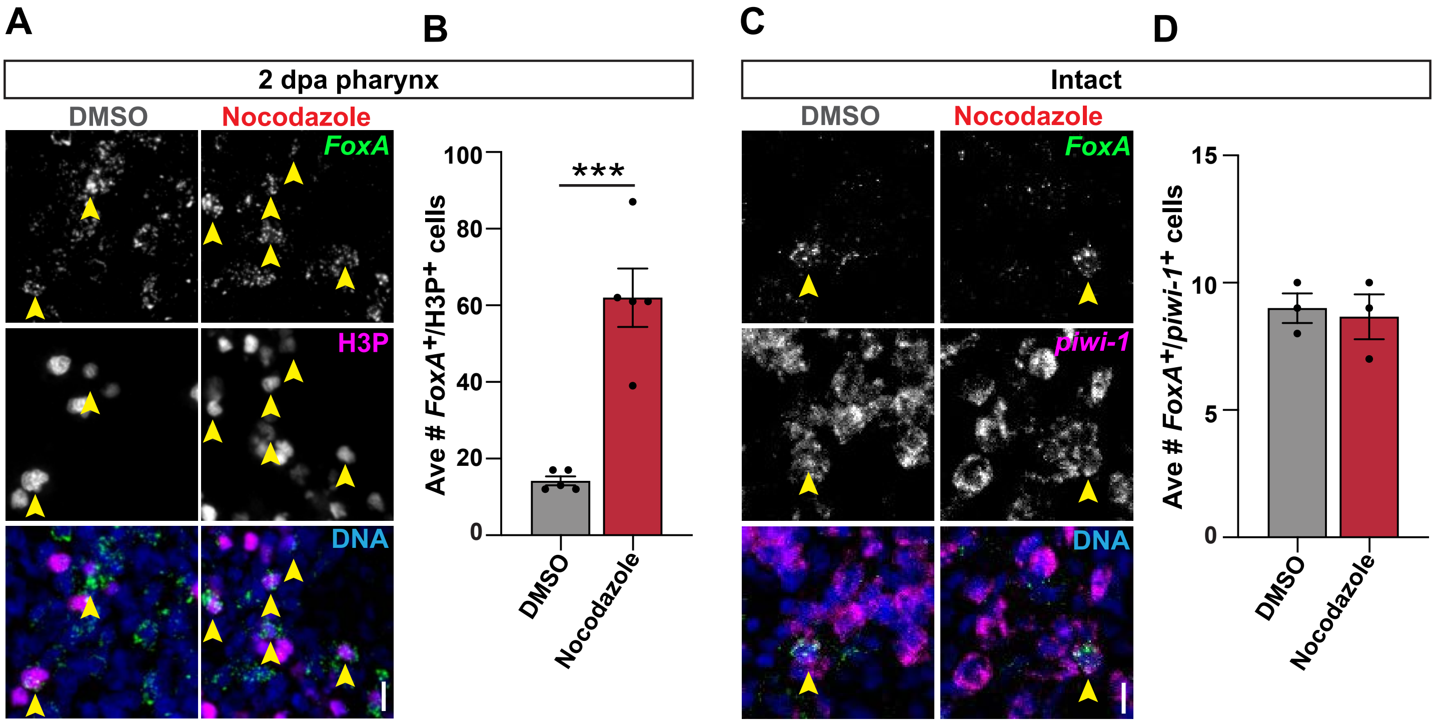

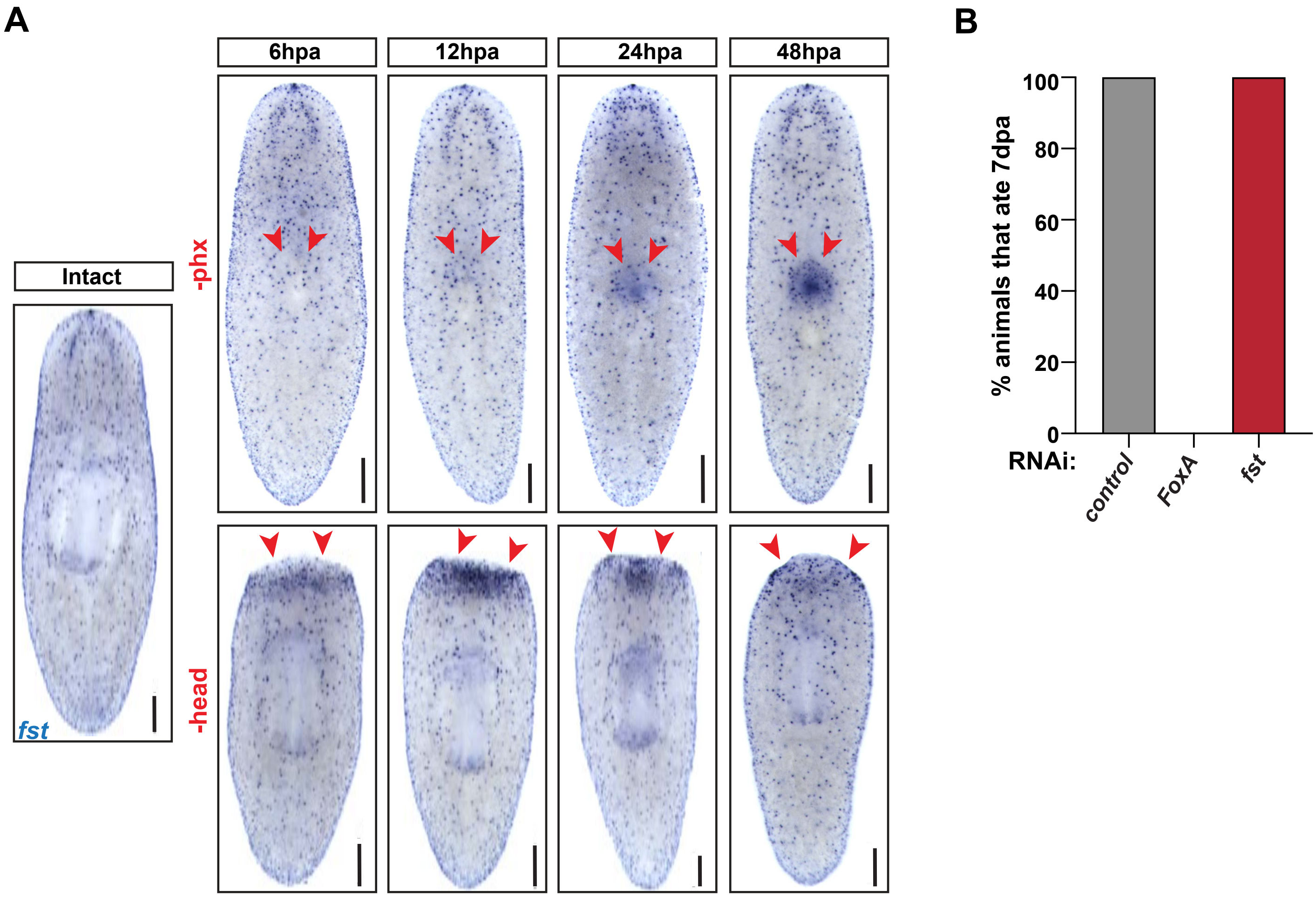

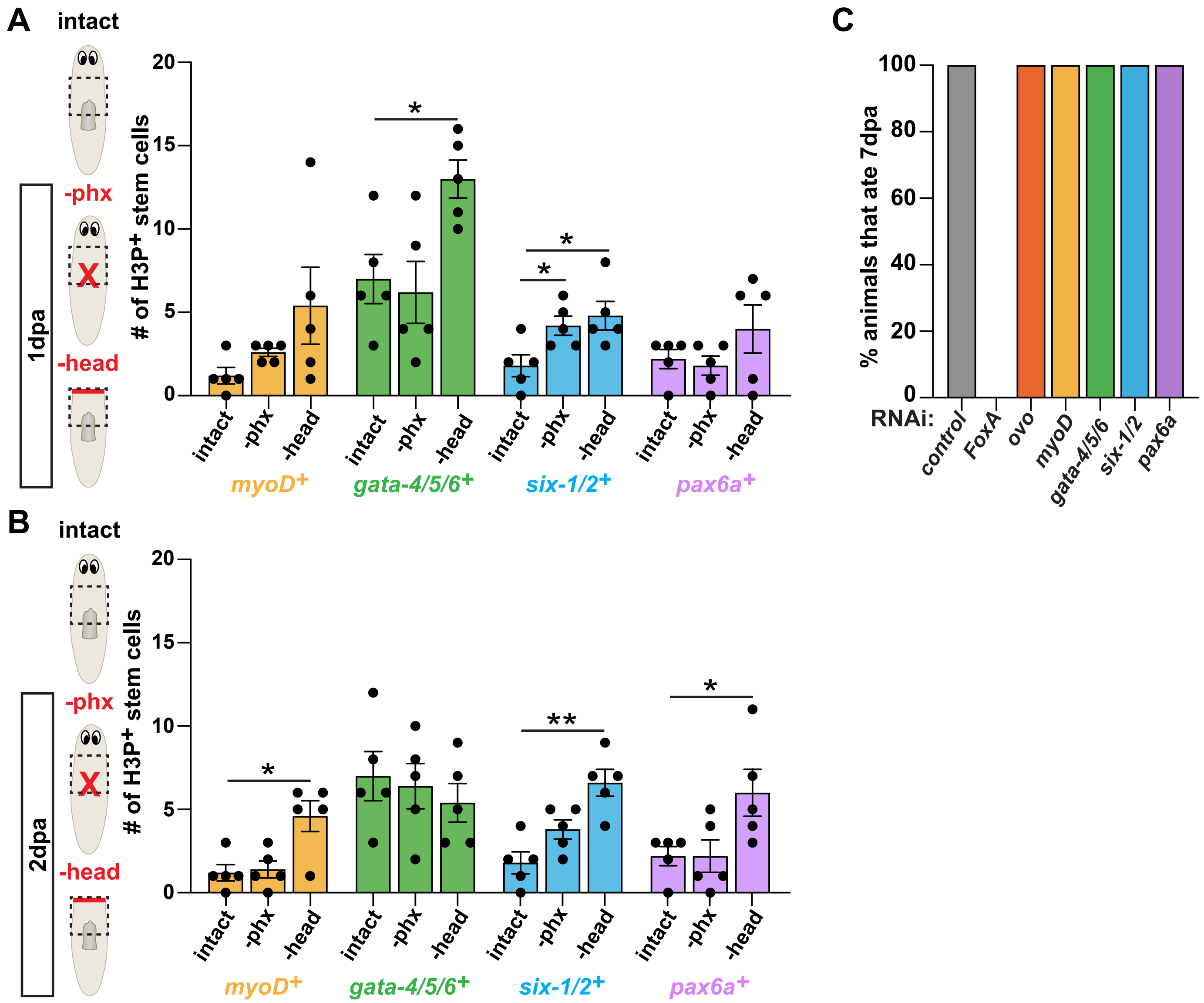
